# Introducing ‘identification probability’ for automated and transferable assessment of metabolite identification confidence in metabolomics and related studies

**DOI:** 10.1101/2024.07.30.605945

**Authors:** Thomas O. Metz, Christine H. Chang, Vasuk Gautam, Afia Anjum, Siyang Tian, Fei Wang, Sean M. Colby, Jamie R. Nunez, Madison R. Blumer, Arthur S. Edison, Oliver Fiehn, Dean P. Jones, Shuzhao Li, Edward T. Morgan, Gary J. Patti, Dylan H. Ross, Madelyn R. Shapiro, Antony J. Williams, David S. Wishart

## Abstract

Methods for assessing compound identification confidence in metabolomics and related studies have been debated and actively researched for the past two decades. The earliest effort in 2007 focused primarily on mass spectrometry and nuclear magnetic resonance spectroscopy and resulted in four recommended levels of metabolite identification confidence – the Metabolite Standards Initiative (MSI) Levels. In 2014, the original MSI Levels were expanded to five levels (including two sublevels) to facilitate communication of compound identification confidence in high resolution mass spectrometry studies. Further refinement in identification levels have occurred, for example to accommodate use of ion mobility spectrometry in metabolomics workflows, and alternate approaches to communicate compound identification confidence also have been developed based on identification points schema. However, neither qualitative levels of identification confidence nor quantitative scoring systems address the degree of ambiguity in compound identifications in context of the chemical space being considered, are easily automated, or are transferable between analytical platforms. In this perspective, we propose that the metabolomics and related communities consider identification probability as an approach for automated and transferable assessment of compound identification and ambiguity in metabolomics and related studies. Identification probability is defined simply as 1/N, where N is the number of compounds in a reference library or chemical space that match to an experimentally measured molecule within user-defined measurement precision(s), for example mass measurement or retention time accuracy, etc. We demonstrate the utility of identification probability in an *in silico* analysis of multi-property reference libraries constructed from the Human Metabolome Database and computational property predictions, provide guidance to the community in transparent implementation of the concept, and invite the community to further evaluate this concept in parallel with their current preferred methods for assessing metabolite identification confidence.

## INTRODUCTION

### Comparing Molecular Identification Among Omics Measurements

In biomedical research, systems biology studies^1–3^ are used to discover new disease biomarkers and elucidate underlying biological mechanisms. Such studies are driven by multiple high-throughput omics technologies: genomics,^4, 5^ transcriptomics,^6^ proteomics^7, 8^ and metabolomics.^9, 10^ Genomics and transcriptomics are the most mature, owing to the more limited chemical diversity of DNA and RNA relative to proteins or metabolites,^11, 12^ the fidelity and accuracy of the associated measurement techniques (i.e. sequencing)^5^, and the breakthrough of having a complete human genome reference sequence as a result of the Human Genome Project.^13^ Today, whole genomes can be sequenced in just 1-2 days with error rates <0.1%,^14^ using modern high throughput sequencing technology (e.g. Illumina NovaSeq) and exploiting the fidelity of DNA polymerase for molecular replication and the specificity of fluorophores read from labeled base pairs.^5^

Proteomics is next in technical maturity. This is because proteins have only slightly greater chemical diversity compared to DNA and RNA, as they are composed of 22 amino acids. However, the complexity of the proteome can increase greatly if all possible protein post-translational modifications (PTMs; e.g. phosphorylation) are considered, and the computational time required for processing mass spectrometry-based proteomics data scales exponentially with the number of PTMs considered. Mass spectrometry-based proteomics^7, 8^ exploits several characteristics of proteins and their constituent peptides. First, proteins are direct readouts of the genetic code, and if the genome is known, then associated protein sequences can be determined.^15^ Second, peptides dissociate characteristically around the amide bond during a tandem mass spectrometry (MS/MS) measurement, allowing for accurate prediction of their fragmentation spectra.^16, 17^ These characteristics have led to analytical workflows that can determine the proteomes of moderately complex samples, as well as methods for estimating and controlling peptide and protein identification false discovery rates (FDRs).^18, 19^ Completely measuring the proteomes of highly complex samples (e.g., human blood plasma) requires a balancing of time and cost. In addition, comprehensive determination of post-translationally modified proteins^20^ and hybrid peptides^21^ remains challenging.

Metabolomics is the least mature among the omics sciences, with high-throughput, untargeted measurements having the goal of identifying and quantifying as many non-protein, small molecules (e.g., 50-1500 Da) as possible. Given their high sensitivity and broad molecular coverage, a variety of liquid chromatography-mass spectrometry (LC-MS) techniques are used in untargeted metabolomics. Typically, LC-MS assays yield thousands of signals with unique *m/z* and retention time (RT) coordinates. Each signal, defined as a “feature”, represents a potential small molecule of interest. However, these features may also be due to chemical noise or contamination and chemical variants of small molecules such as protonated- or sodiated-adducts. The rate-limiting step in untargeted metabolomics is discerning among these signals to annotate the chemical structures associated with the detected features. The current paradigm for confident metabolite identification involves comparing experimental MS (or nuclear magnetic resonance spectroscopy; NMR) data from biological measurements to comparable data from purified reference metabolites that were measured under similar conditions, preferably in the same laboratory. Unlike proteomics, where the analytes of interest are encoded by the genome and limited to linear polymers of repeating amino acids, the chemical space being profiled in metabolomics is essentially unconstrained, especially if exogenous metabolites (such as food products), microbial transformations and other chemical exposures are considered. As a result, much less is known about the complete composition of the human metabolome than the genome or the proteome. This is because relatively few reference standards are available relative to the known chemical space, and the measured properties such as mass fragmentation patterns are less predictable for metabolites than peptides. Consequently, metabolite identification in metabolomics is often prone to more errors or uncertainties than other omics technologies. Even if we could computationally predict all metabolites likely to exist in a given organism or biofluid, based on genomes or proteomes, the search would not be complete due to interaction of the organism with non-biological sources. As a result, even though metabolomics reference libraries continue to grow^22, 23^, they are unlikely to ever be complete. This has inspired efforts to increase reference data through enzymatic biotransformation of drugs and other xenobiotic chemicals.^24^

### Placing Confidence in Metabolomics Identifications

Insufficient knowledge of, or constraints placed upon, which small molecules might be present in a sample creates unique challenges when attempting to identify the chemical structure associated with a feature detected in metabolomics analyses. Even with the most recent developments in software and innovative computational methods that can automate steps in the informatics workflow,^25, 26^ a critical question is the level of confidence that one has in the identifications proposed. The extent to which experimental data collected from a sample matches reference data is typically used to support feature identification. Unfortunately, there are often dozens to thousands of possible isomers in chemical and metabolomic databases. Isomers are chemicals with the same elemental formulae but different three-dimensional structures or different atomic positions. MS alone is not capable of disambiguating most of these isomers. In addition, other kinds of isomers may exist for which reference data do not yet exist.^27^ Hence, apart from the accurate mass (monoisotopic *m/z*) data and MS/MS spectra obtained from a MS measurement, complementary data from additional analytical measurements (e.g., retention time, collision cross section (CCS), NMR spectra, different ionization modes or chemical derivatizations) improve identification confidence by limiting the number of potential compounds that satisfy the given match criteria^28^. However, currently there is no method for quantifying the ambiguity in a metabolite identification in context of the chemical space being considered. Accurately estimating total FDR in compound identifications is still in its infancy in metabolomics.^29–31^

In 2005, a Metabolomics Standards Workshop^32^ was convened by the U.S. National Institutes of Health and the Metabolomics Society with the goal of establishing a Metabolomics Standards Initiative (MSI)^33^ that would consider and recommend minimum reporting standards for describing various aspects of metabolomics experiments. The MSI consisted of five working groups comprised of international experts in metabolomics research and that developed recommended requirements for biological context, chemical analysis, data processing, ontology, and data exchange associated with metabolomics studies. In 2007, the Chemical Analysis Working Group of the MSI published the seminal paper on the minimum information for reporting the chemical analysis metadata associated with a metabolomics study, including a 4-level, qualitative scheme for reporting metabolite identification confidence.^28^ These MSI-levels have been revised to include additional considerations^34^ or other data types^35^ but have remained largely unchanged. In 2014, Sumner *et al.*^36^ and Creek *et al.*^37^ proposed a transition from the existing qualitative metabolite identification confidence levels to a quantitative scoring system based on identification points (IP), citing the bias of the traditional MSI-levels towards identifications made in the context of data from authentic reference compounds or the need for more granularity in the levels, respectively. Most recently, Alygizakis and colleagues used a machine learning approach to develop a new IP-based system.^38^ All three papers cited the EU Guideline 2002/657/EC^39^ as a motivating example.

Reporting qualitative MSI-confidence levels in metabolite identifications is infrequently and inconsistently used by members of the metabolomics community. This is likely because assigning confidence scores is still a subjective process for most data reporters. Recipients of such data reports lack sufficient information or tools to independently verify metabolite identifications. Many reports include only chemical names, but not chemical or structure identifiers like PubChem Compound Identifications (PubChem CIDs) or International Chemical Identifiers (InChIs).^40^ Chemical names can be highly ambiguous and misleading for data consumers and easily lead to problems in comparing data across different biological studies, as recently highlighted by arguments in the lipidomics literature.^41^ For scientists who process LC-MS/MS data, deciding whether a given experimental MS/MS spectrum matches a reference spectrum is dependent on the metric used, the threshold set, and many other ambiguous decision points.^29^ The current best alternative to MSI-confidence levels is to provide both raw and processed data in public repositories such as the Metabolomics Workbench^42^ and MetaboLights^43^ to support claims of reported metabolite identifications and to allow for independent verification.

### Expanding from Measures of Confidence to Measures of Ambiguity

For both qualitative levels of metabolite identification confidence^28, 34, 35^ and quantitative scoring systems^36, 37^, the methods are not easily automated or transferable between analytical platforms (e.g., MS and NMR) and the degree of ambiguity or uncertainty in identifications is not fully represented. That is, given a reference library of a certain size and composition, and an analytical approach of certain resolution and precision, what is the likelihood of one identification being more correct than another given the available evidence? Here, we introduce a concept for moving from levels of identification confidence or cumulative point scoring systems to a universal method that assigns a mathematical probability to a given identification being correct. Importantly, this concept considers the composition and size of the reference library used, the numbers and types of measurement dimensions included in the experimental analysis, and each measurement’s precision. It is also easily automated and the results transferable between analytical platforms.

## INTRODUCTION TO METABOLITE IDENTIFICATION PROBABILITY

### Logic Supporting the Concept

Metabolite identification probability represents a first step in moving away from semi-manually assigning subjective, constantly evolving levels of identification confidence (e.g., MSI levels) or IP-based methods towards a universal, automated method. Importantly, while methods for estimating FDRs for non-peptide small molecules have been explored in the context of MS/MS spectral matching, these have not been extended to other technologies (e.g., NMR) and data types (e.g., retention times, CCS values). The identification probability concept that we introduce here can be applied to any metabolomics measurement technology or method that relies on reference libraries (e.g., MS, GC-MS, LC-MS, LC-ion mobility spectrometry (IMS), LC-IMS-MS, LC-IMS-MS/MS, NMR, LC-NMR, etc.). Identification probability is defined as follows:

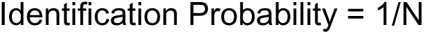

where N is the number of molecules in a reference library that match an experimentally measured feature within the precision(s) of the given measurement technology or method and the user-defined tolerances allowed in the measurement precision(s)

Based on this definition, higher dimensional analytical approaches or those that provide measurements of more properties should provide higher probability in a compound identification due to their ability to provide higher resolution of chemical space, while larger reference libraries would make it more difficult to completely resolve molecules in chemical space due to higher potential for conflicts.

Let’s consider a single dimension or single property analysis to start. MS when used alone produces mass spectra, and the spectra will have a given resolution, based on the type of mass spectrometer used. Fourier transform ion cyclotron resonance (FTICR)-MS provides the highest mass resolution among current mass spectrometers used for metabolomics and related studies and can lead to extremely high accuracy in determining the exact molecular formulae that correspond to detected isotope patterns in the mass spectrum. The determined molecular formulae can then be searched against an appropriate reference library consisting of known molecular formulae; in our example, we consider a subset of the Human Metabolome Database (HMDB)^44^ consisting of 22,077 non-lipid molecules for which computationally-predicted reference data were generated (**Supplemental Table S1**). If one were to perform a metabolomics experiment and detect a feature with protonated exact mass equal to 116.07115 Daltons, then the calculated molecular formula would be C5H9NO2, which <u>may</u> correspond to the target molecule 4-amino-2-methylenebutanoic acid. When that formula is searched against the down-selected HMDB library that contains molecular formulae for up to 9 compounds with the same formula, then the probability of the experimentally measured formula C5H9NO2 actually being 4-amino-2-methylenebutanoic acid (or any of the 9 candidates) would be 1/9 or 11% (**Figure 1**). Now, on the other end of the extreme, let’s consider a multi-dimensional analysis, such as IMS-MS/MS. From this analysis, we would determine an IMS drift time or CCS value, a MS/MS spectrum and an accurate mass. The individual measurement precisions of any of these dimensions is not sufficiently high as to allow exact determination of any given property, and so matching of experimental data to the library proceeds within ranges or tolerances determined by typical experimental precision: ± 10 ppm for mass, ± 1% for CCS, and ≥ 850 for cosine similarity score (for MS/MS spectral matching). In the example shown in **Figure 1** for the target molecule 4-amino-2-methylenebutanoic acid, the combination of ± 10 ppm and ± 1% CCS reduces the candidates in the reference library to 7, and the identification probability for all candidates is 1/7 or 14%. For the same example, the combination of ± 10 ppm, ± 1% CCS, and ≥ 850 cosine similarity score reduces the candidates in the reference library to 1, and the identification probability is 1/1 or 100% for the measured feature corresponding to the target molecule 4-amino-2-methylenebutanoic acid. A key advantage to higher dimensional analyses is that the likelihood in complete overlap among property sets for library entries decreases roughly in proportion to the number of dimensions of the analysis.

**Figure 1.**
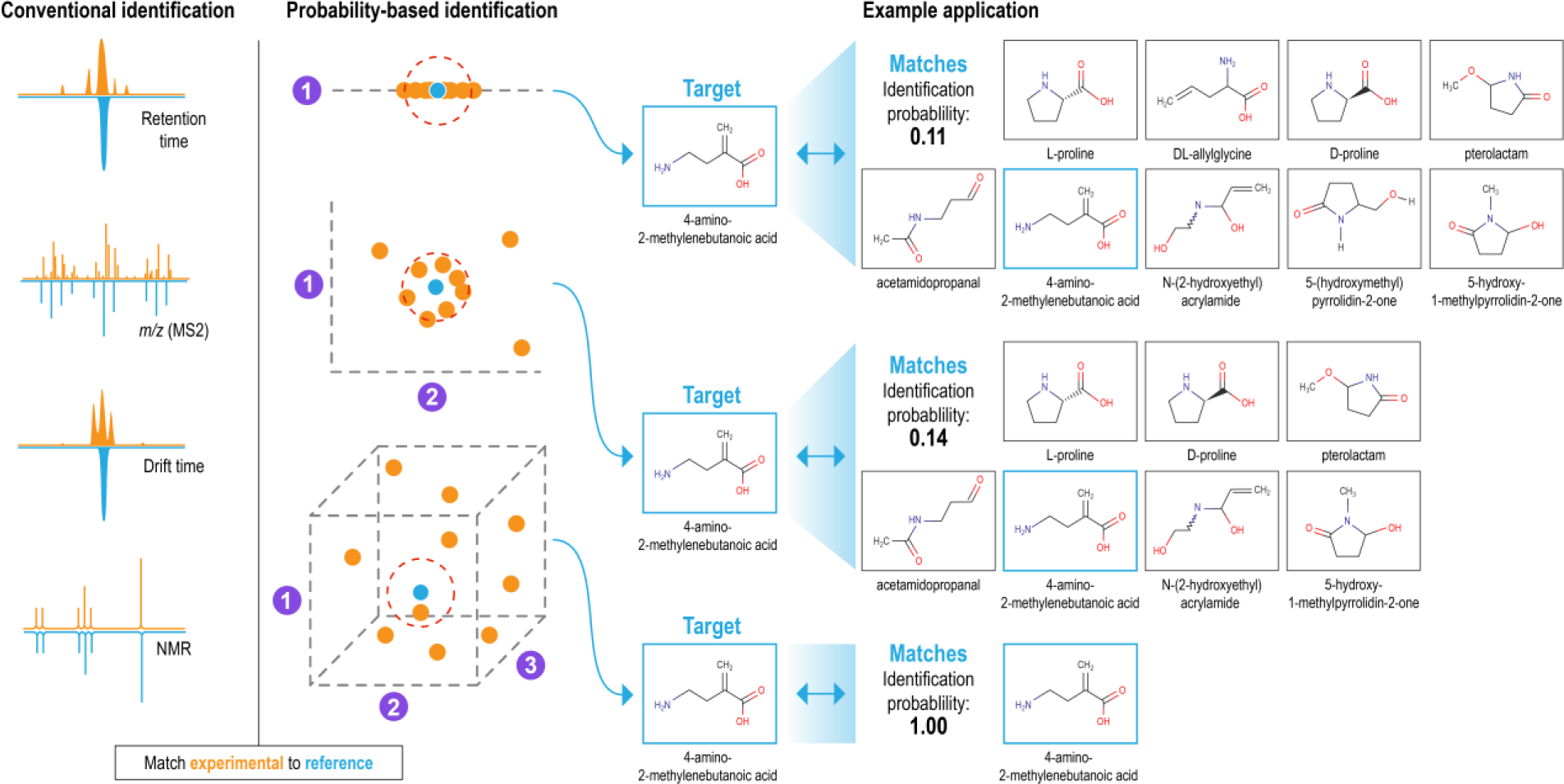
Demonstration of the metabolite identification probability concept using 4-amino-2-methylenebutanoic acid as the target molecule. Conventional metabolite identification (left panel) is based on manual or semi-automated comparison of experimental data to similar data contained in reference libraries, with final identification confidence determined by a data analyst. Probability-based identification (right panel) is similarly based on comparison of experimental and reference library data and is fully automated. Identification probability is defined as 1/N, where N is the number of molecules in the reference library that match an experimentally measured feature within the precision(s) of the given measurement technology or method and the user-defined tolerances allowed in the measurement precision(s). In the examples shown, the target molecule is 4-amino-2-methylenebutanoic acid, and the reference library is a subset of HMDB consisting of 22,007 non-lipid molecules. In the top row, identification is based on a single dimension of analysis, formula match (1). In the middle row, identification is based on the combination of ± 10 ppm and ± 1% CCS matching (2). In the bottom row, identification is based on the combination of ± 10 ppm, ± 1% CCS, and ≥ 850 cosine similarity score match (3).

### Impacts of reference library size, property match tolerances, and analysis dimensionality

To evaluate how library size, property match tolerances, and dimensionality of analytical analysis might impact metabolite identification probabilities, we further explored the 22,077 non-lipid molecules from HMDB, as well as a complementary set of 44,537 lipid molecules (**Supplemental Table S2**), from the same source. The molecules were classified into a chemical ontology using the ClassyFire tool,^45^ and compounds with an invalid chemical classification value (“NA”) were excluded (**Supplemental Figure S1**). The two molecule sets were placed in separate matrices, together with the protonated mass (*m/z*), RT, CCS, and MS/MS spectra for each molecule. The protonated mass was calculated from the protonated molecular formula. DarkChem^46^ was used to predict CCS for all lipid molecules; for non-lipid molecules, DarkChem, AllCCS,^47^ and DeepCCS^48^ were used to predict CCS for 9308, 10,669, and 2100 molecules, respectively, as indicated in **Supplemental Table S1**. RTs were predicted using Retip^49^ under hydrophilic interaction liquid chromatography (HILIC) conditions for non-lipid molecules or reversed-phase chromatography conditions for lipid molecules (based on ClassyFire assigned superclass of “Lipids or Lipid-like molecules”), and MS/MS spectra were predicted using CFM-ID 4.0^50^ at a “medium” collision energy level of 20 eV. We then matched each of the two molecule sets and their calculated/predicted properties to themselves to simulate the processing of a metabolomics data set.

#### Impact of reference library size

To evaluate the impact of reference library size on metabolite identification probability, we performed Monte Carlo simulations to randomly draw smaller library subsets (e.g. 1,000, 5,000, or 10,000 molecules) from the full lipids and non-lipids libraries. We evaluated 100 randomly drawn subsets for each library size, matched each subset to itself by mass (±10 ppm), aggregated results, and compared the number of matches returned per database search (Error! Reference source not found.**2**). Our results demonstrate that as the size o f the library increases, the relative proportion of matches at a given probability decreases; thus, smaller reference libraries will tend to yield artificially high identification probabilities. Comparing lipids vs non-lipids, the impact of reference library size on identification probability is more pronounced for libraries with more heterogenous content.

**Figure 2.**
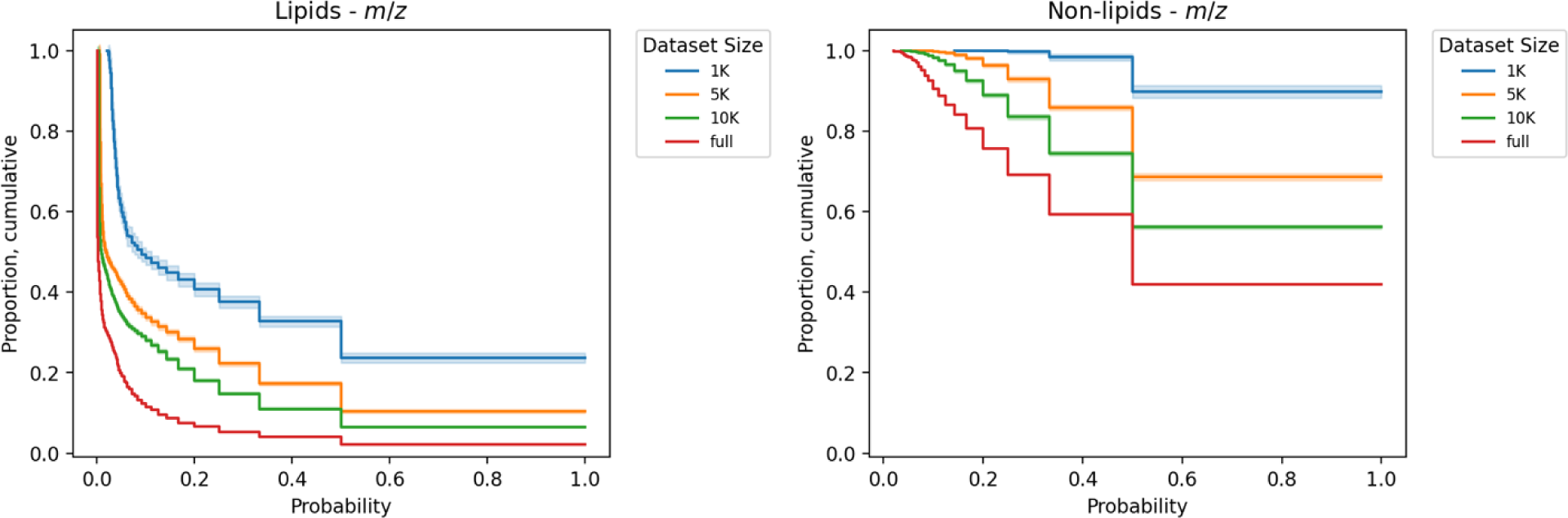
Impact of reference library size on metabolite identification probability. Monte Carlo simulations were performed to randomly draw subsets of the full lipids (left panel) and non-lipids (right panel) reference libraries of size 1K, 5K, and 10K. Match probability is shown on the x-axis, and the proportion of compounds in each dataset matched within ±10 ppm and with a given probability is shown on the y-axis. For example, for non-lipids, a little over 40% of compounds are unambiguously matched when matching the full library to itself with a mass tolerance of ± 10 ppm. Solid lines indicate the mean value, and shaded regions indicate ± 1 standard deviation from the mean based on 100 Monte Carlo simulations (note that no shaded region exists for the full dataset, for which random subsets were not drawn).

#### Impact of property match tolerances and dimensionality of analytical analysis

We next evaluated the impacts of individual property match tolerances and the dimensionality of the analytical analysis on metabolite identification probability. Overall, varying property match tolerance has different impacts on the number of unambiguous identifications depending on the property considered. For instance, the evaluated *m/z* match thresholds gave rise to little, if any, change in the proportion of unambiguous matches from both the lipids and non-lipids datasets, either alone or in combination with other properties (Error! Reference source not found.**3**). We hypothesize that the low variance in match performance across *m/z* tolerances can be attributed to the relative density of compounds occupying *m/z* space vs. the variability of the error thresholds in practical terms. For instance, at an *m/z* of 800 Da (close to the median *m/z* for lipids of 821.8 Da), the error thresholds of ± 0.1, 1, 5, and 10 ppm correspond to ± 0.00008 Da, ± 0.0008 Da, ± 0.004 Da, and ± 0.008 Da, respectively. The resolutions may not differ sufficiently to effect significant changes to the number of matches within each corresponding tolerance.

In contrast to *m/z*, CCS search tolerance has a more pronounced impact on unambiguous matches. While searching by CCS alone produces zero or near-zero unambiguous matches across both lipids and non-lipids datasets, when used in combination with other analytical dimensions, the effect of CCS search tolerance becomes much more pronounced. In some cases, we observe a two-fold or even greater increase in the fraction of the dataset which can be definitively matched, particularly in the case of lipids (Error! Reference source not found.**3**). Our simulation data suggests that when used in conjunction with other measurements, accurate CCS measurements have the potential to increase the number of confident identifications. However, we note that the highest-accuracy CCS error threshold evaluated is a CCS error of ± 0.1%, which may be achievable experimentally only using very high-resolution ion mobility separations, such as structures for lossless ion manipulations (SLIM).^51^ Inclusion of CCS in compound matching at this tighter threshold produced marked improvements in the proportion of unambiguous matches.

Neither of the two RT thresholds evaluated (± 0.1 min and ± 0.5 min) produced any unambiguous matches in the database using RT alone for the lipids or non-lipids datasets (data not shown). However, as with CCS, RT combined with additional measurement dimensions produced more confident matches (**Figure 3**). Reducing the RT tolerance from 0.5 min to 0.1 min correspondingly increases the proportion of unambiguous matches. While the observed effect is smaller than the impact of CCS, the inclusion of RT still substantially improves unambiguous identifications, especially compared with *m/z*.

**Figure 3.**
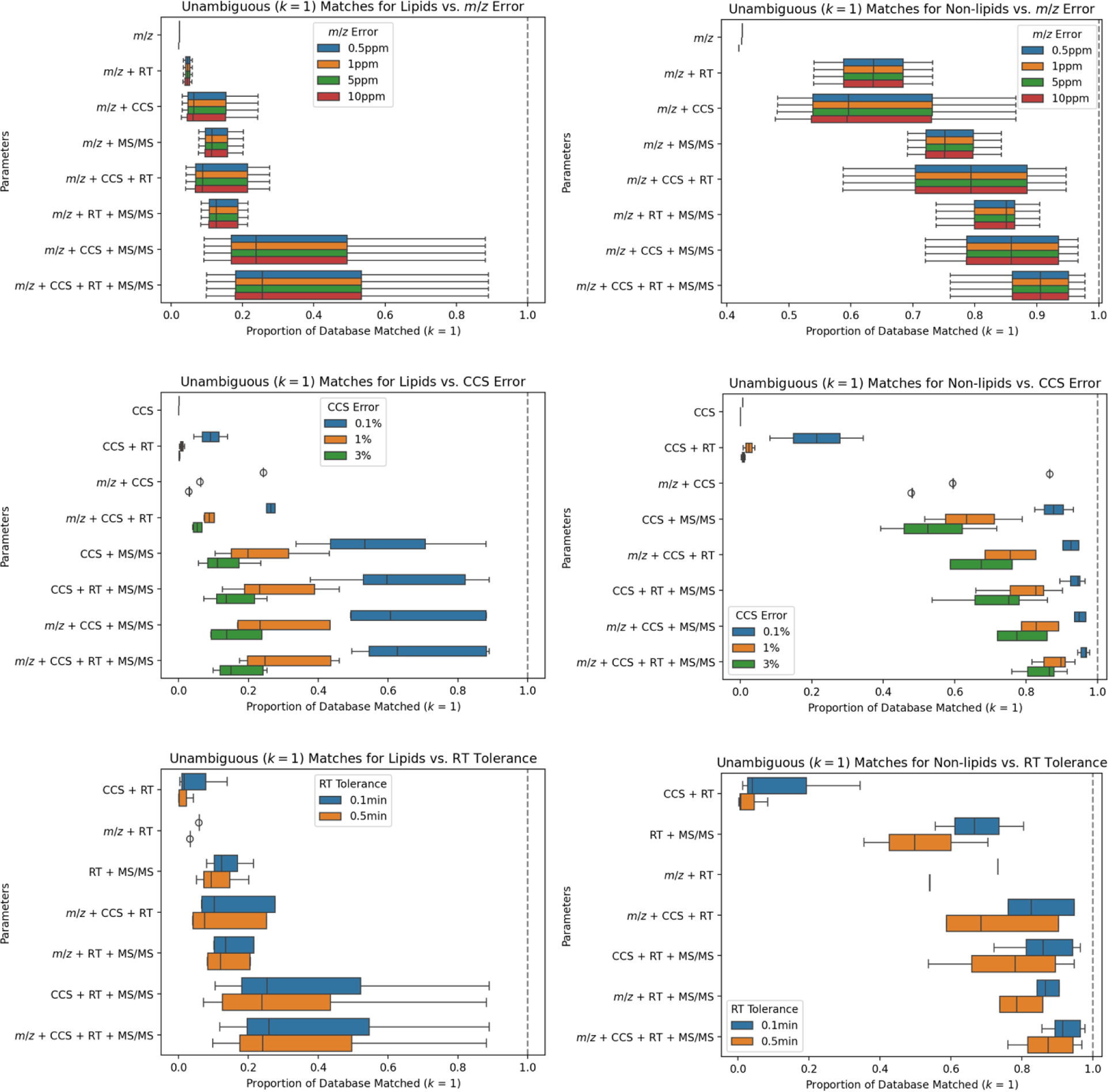
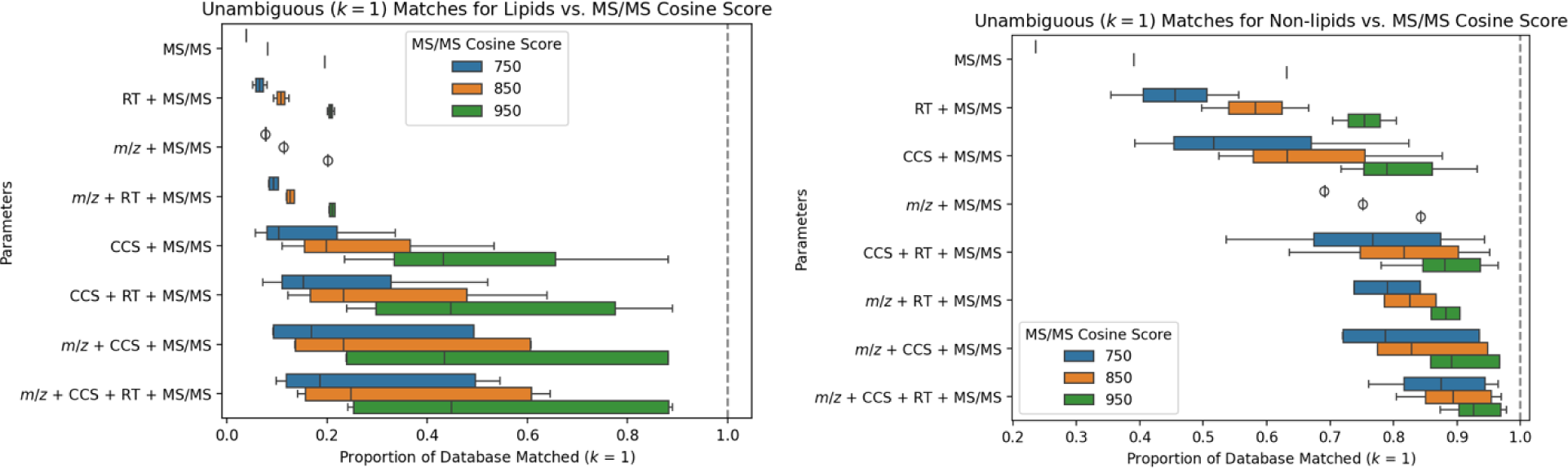
Impact of property match tolerances and dimensionality of analytical analysis on metabolite identification probability for lipids (left) and non-lipids (right). Each boxplot summarizes the fraction of each database that is unambiguously matched (*k* = 1) when varying the search tolerances evaluated in each dimension (*m/z*, CCS, RT, and MS/MS, respectively) as shown. The first set of boxplots in each plot represent results when only considering the dimension of interest and varying search tolerance within that single dimension, with the subsequent boxplots depicting results upon inclusion of additional search dimensions but only varying the search tolerance of the first dimension. For each dimension, search tolerances include *m/z* ± 0.1 ppm, ± 1 ppm, ± 5 ppm, and ± 10 ppm; CCS ± 0.1%, ± 1%, and ± 3%; RT ± 0.1 min, and ± 0.5 min; and MS/MS cosine score ≥ 750, ≥ 850, and ≥ 950.

Finally, we evaluated the impact of the MS/MS spectral match threshold. We chose to use cosine similarity score due to its ubiquitous use; however, we note that alternative scoring algorithms, such as spectral entropy,^29^ have demonstrated improvements over cosine similarity. Based on the range of typical scoring thresholds used for MS/MS matching, we evaluated cosine similarity thresholds of 750, 850, and 950. Our results show that among both lipids and non-lipids, MS/MS score alone is the best-performing singular measurement in terms of identifying compounds unambiguously (**Figure 3**). In contrast to *m/z*, however, increasing the MS/MS cosine score threshold resulted in significant increases to the proportion of compounds producing unambiguous matches in both the lipids and non-lipids libraries. In fact, when matching by MS/MS cosine score alone, 63% of non-lipids can be accurately matched with a cosine similarity score of ≥950, compared to just 24% with a cosine score of 750. As before, the MS/MS dimension can be combined with other measurement dimensions to achieve an even greater fraction of unambiguous identifications; in fact, all the best-performing multi-dimensional search parameter sets include MS/MS.

While the example data and toy metabolite identification probability analyses discussed above are LC-MS-centric, the concept is applicable for any workflow that produces metabolite identifications through matching experimental data to similar data in reference libraries, such as NMR and GC-MS. Indeed, many NMR spectral matching algorithms, such as those used in MagMet^52^, Bayesil^53^ and Chenomx^54^, use concepts similar to the cosine similarity score used in MS/MS. Likewise, GC-MS uses equivalent concepts as LC-MS/MS for spectral matching.

## THE ROLE OF REFERENCE LIBRARIES AND HOW TO POPULATE THEM

### Current Landscape and Use of Reference Libraries for Compound Identification

The metabolite identification probability concept introduced here depends on the size and contents of the reference library used. As such, it is essential that reference libraries are populated and used correctly. In the following discussion, we describe the current landscape and use of reference libraries for compound identification and provide recommendations to the community for their use as it relates to metabolite identification probability. For the purposes of this discussion, we assume that the contents of these reference libraries are correct and accurate.

Reference libraries contain varying levels of curated information about compounds (e.g., structure, properties, and classifications). At a minimum, useful reference libraries contain compound structures in machine readable formats or public identifiers that map to chemical structures, alongside derived properties such as elemental formulae and exact monoisotopic masses. In particular, many reference libraries developed for use with specific analytical approaches contain measurable observables, such as observed precursor ions and MS/MS or NMR spectra. They also include experimental metadata that define these spectra, such as the type of instrument used or e.g., details of the MS/MS fragmentation method that was applied. For the case of high-resolution MS (HRMS), the data can be used to directly search against exact masses of known, expected, and even predicted chemical structures. If HRMS data accuracy of <0.002 Da is achieved, chemical formulae can be inferred using a variety of different software tools, especially if MS/MS and isotope ratio information is included.^55–59^ Note that a mass resolving power of R<250,000 means that alternative formulae might still need to be considered.^60^

Many open-access reference libraries exist in the form of compound collections that contain mass, formula and structure information for millions of known, suspected or predicted compounds (**Table 1**). These include PubChem^61^ which has nearly 110 million compounds, ChEMBL^62^ with 2.1 million compounds, and the US-EPA CompTox Chemicals Dashboard^63^ with 1.2 million compounds. All of these support mass and formula searching. However, they also include a large fraction (>99%) of anthropogenic molecules, making these libraries somewhat more suited for exposomics^64^ or environmental testing studies and less suitable for traditional metabolomics studies that focus on physiological metabolites. A number of reference libraries exist that focus on storing only known biologically-related compounds. For example, the Human Metabolome Database (HMDB) now accounts for 248,097 compounds,^44^ Lipid Maps^65^ lists 45,684 compounds, KEGG^66^ denotes 18,784 compounds, and MetaCyc^67^ includes 16,861 compounds. These databases continue to expand in coverage and content and such databases are much more suitable for traditional metabolomics studies.

**Table 1.** Compound collection reference libraries. These reference libraries function primarily as collections of compounds and include chemical structures, molecular formulae, masses and physicochemical properties, among other data.

| Library | Number of Compounds | URL | Citation |
| --- | --- | --- | --- |
| ChemSpider | >129,000,000 | http://www.chemspider.com/ | 68 |
| PubChem | >119,000,000 | https://pubchem.ncbi.nlm.nih.gov/ | 69 |
| CompTox Chemicals Dashboard | >1,200,000 | https://comptox.epa.gov/dashboard/ | 63 |
| RaMP-DB 2.0 | >256,000 | https://rampdb.nih.gov/ | 70 |
| Human Metabolome Database (HMDB) | >250,000 | https://hmdb.ca/ | 44 |
| Metabolomics Workbench | >164,000 | https://www.metabolomicsworkbench.org/ | 42 |
| Chemical Entities of Biological Interest (ChEBI) | >160,000 | https://www.ebi.ac.uk/chebi/ | 71 |
| LipidMaps | >47,000 | https://www.lipidmaps.org/databases/lmsd/overview | 65 |
| Natural Products Atlas | >33,000 | https://www.npatlas.org/ | 72 |
| Metabolights | >27,000 | https://www.ebi.ac.uk/metabolights/index | 43 |
| Kyoto Encyclopedia of Genes and Genomes (KEGG) | >19,000 | https://www.genome.jp/kegg/ | 66 |
| MetaCyc | >18,000 | https://metacyc.org/ | 67 |

While *m/z* or formula searching is relatively easy to perform, and the sizes of the reference libraries mentioned above are often very large, the reliability of these single parameter matches is often quite poor. Indeed, it is often possible to get hundreds of potential matches with a single *m/z,* or even formula, query (**Figure 4**).^73, 74^ Additional “observable” information is needed to add specificity and increase confidence in tentative compound identifications.^28, 34^ Generally, the most accessible and reproducible experimental measurements, beyond molecular weight, are spectral or separations data. This includes MS/MS spectra (for LC-MS or CE-MS), electron ionization (EI) spectra (for GC-MS) or NMR spectra, and retention times (RT; for LC) or retention indices (RI; for GC) and drift times or collision cross sections (CCS) for IMS data. More recent technology developments allow for the collection of infrared spectra in-line with IMS and MS measurements.^75^ The intensity, position, number and character of the peaks seen in MS/MS or NMR spectra is often considered sufficient to make identifications of metabolites; however, as shown in **Figure 3**, MS/MS spectral match alone is insufficient for providing unambiguous identification of metabolites when matching to large reference libraries. Several different scoring schemes are available to facilitate spectral matching and scoring and offer superior results to simply matching based on a mass or formula.^76, 77^ Recently, spectral entropy was developed as a new MS/MS scoring scheme to particularly account for spectra with few fragment ions, as often observed in small molecule analyses.^29^ The chromatographic and separation parameters are related to physicochemical properties (e.g., size, shape, charge, boiling point, hydrophobicity) and provide information that is fundamentally different from measured mass or fragmentation spectra. RI and CCS values can be relatively instrument- or condition-independent with proper calibration, making them highly reproducible and suitable for compound identification. CCS values are particularly reproducible, with relative standard deviations <1% reported in interlaboratory comparisons and under standardized conditions.^78^ Fragmentation spectra (from GC-MS or LC-MS/MS) are generally relied upon the most in identification workflows due to their specificity and wide availability of associated instrumentation. GC-electron ionization mass spectra were standardized over 60 years ago. Yet, in comparison, measured spectra from LC-MS/MS are harder to standardize due to the variability between instruments, the fragmentation conditions and the collision energies used. Therefore, MS/MS libraries often contain multiple spectra for each compound.

**Figure 4.**
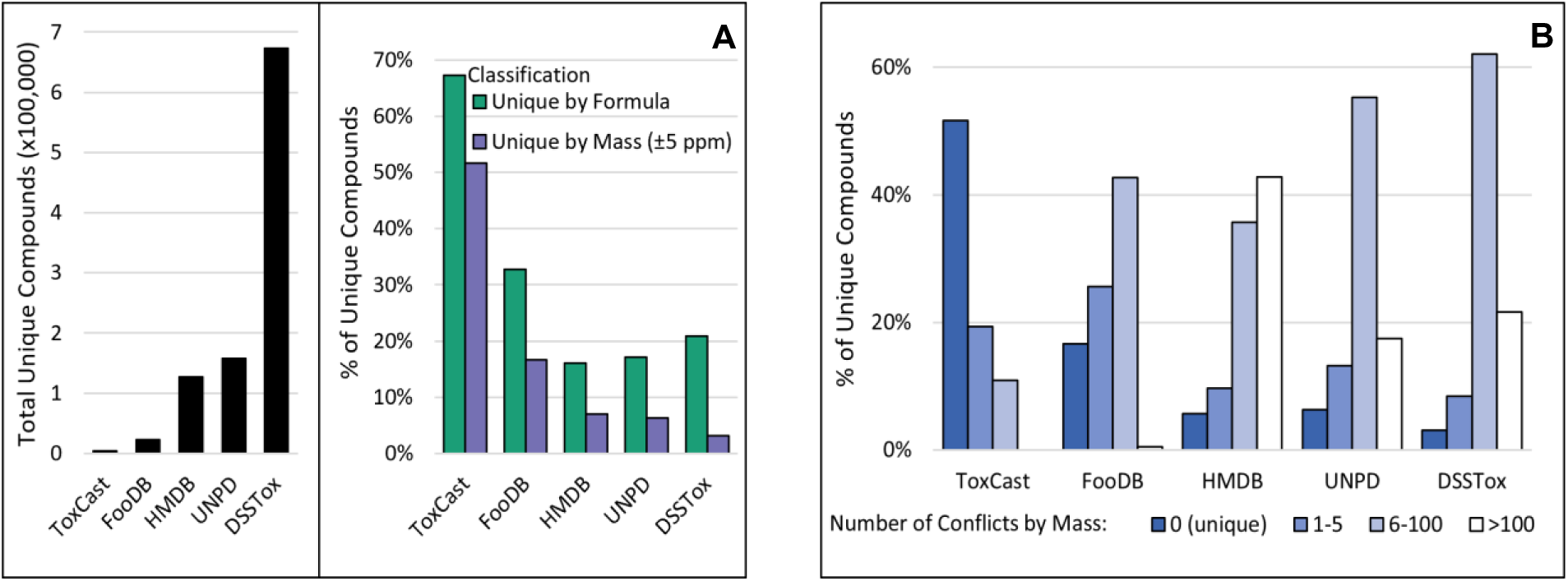
Size, composition, and uniqueness of representative reference libraries. (A) Number of unique compounds by structure (based on unique canonical SMILES generated by RDKit; Left Panel), and percent of compounds unique by formula or parent mass (Right Panel). Unique by parent mass indicates that there are no other compounds with a mass within 5 ppm. (B) Percent of library compounds that conflict by parent mass with up to >100 other molecules. Number of conflicts by parent mass is a count for how many other structures within the given library have a mass within 5 ppm. If there are 0 conflicts for a given structure, it is considered unique by parent mass.

Because of their utility in providing additional confidence in metabolite identification, there are a growing number of both commercial and open-access reference libraries that contain various properties from experimental measurements of pure reference compounds and that are available for matching to metabolomics data. Popular reference libraries that contain mass spectral data are MassBank.eu, MassBank of North America (MassBank.us), the NIST spectral library,^77^ METLIN,^22^ and mzCloud, as well as commercial libraries produced by Waters, Sciex, Bruker, Agilent, and Thermo Fisher. Other resources exist that contain both spectra from analysis of pure compounds but also large numbers of spectra of unknown compounds from analysis of real samples, such as GNPS.^79^ Some of the more popular NMR spectral libraries are the BioMagResBank,^80^ NMRShiftDB,^81^ NP-MRD,^82^ and COLMAR,^83^ as well as commercial libraries produced by Bruker and Chenomx. Popular reference libraries that contain RI and/or CCS include: the NIST RI library, the FiehnLib RI library,^84^ the Unified CCS Compendium,^85^ the Sumner CCS library^86^ and several commercial CCS libraries from instrument vendors such as Bruker, Agilent and Waters. MassBank.us contains many metabolites with LC-based retention times, including for hydrophilic interaction chromatography (HILIC)^49^. In contrast to standardized gas chromatography RI and CCS measurements, LC RT and electrophoretic mobilities are not easily translated from instrument to instrument or from one configuration to another. As a result, reference libraries for LC RT and electrophoretic mobility are often quite small. Recently however, the developers of METLIN released a reference library containing >80,000 RTs measured for small molecules, called SMRT.^87^ These data, the largest of their kind, were collected using a single standard chromatographic protocol but has not been validated yet by independent means. A more detailed listing of reference libraries focused on housing data from analyses of pure reference compounds, their contents, and the number of entries found is provided in **Table 2**.

**Table 2.**
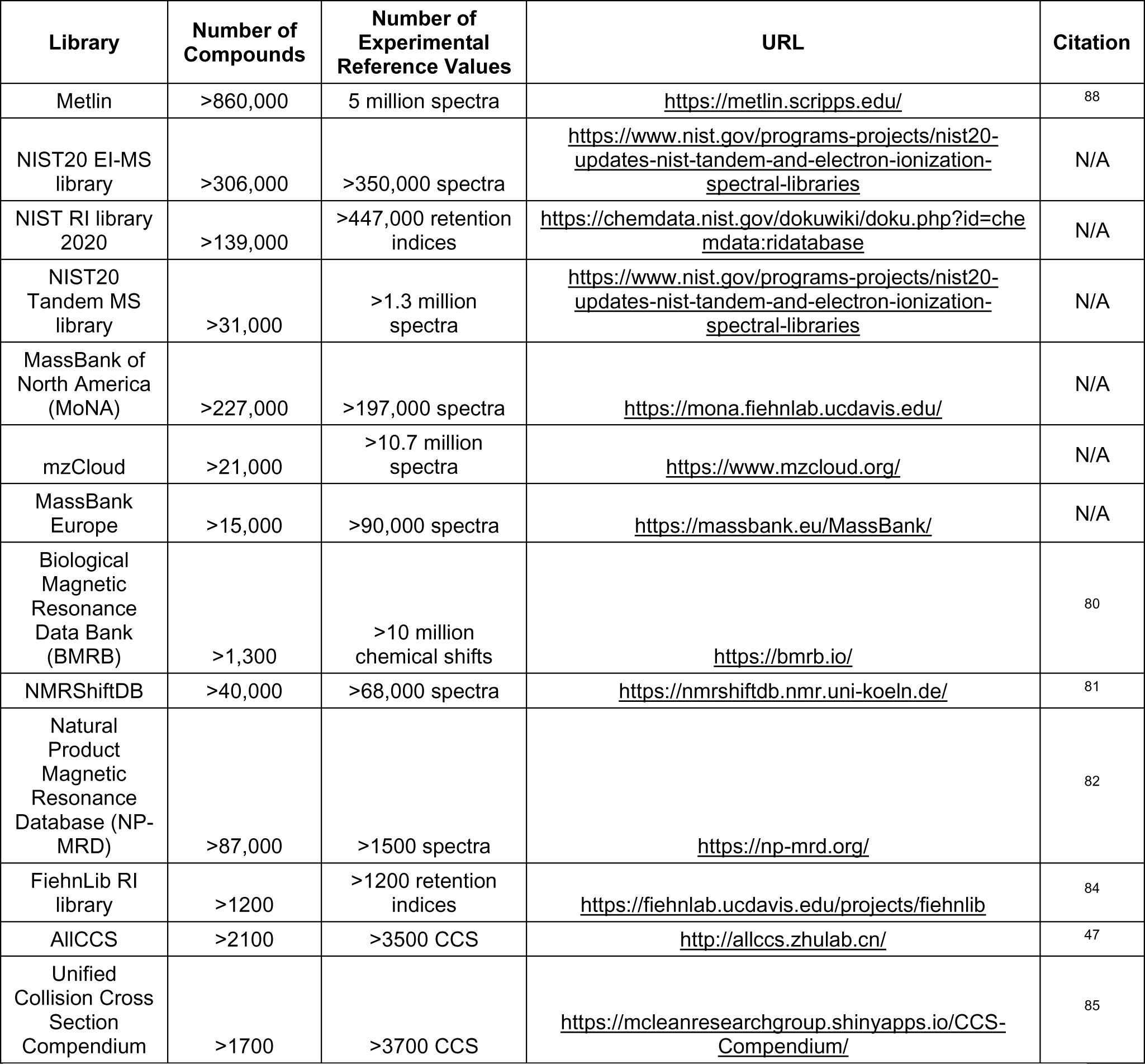
Reference libraries of observable data. These reference libraries contain listings of compounds and their observable data, such as mass spectra, retention indices, NMR spectra and CCS values.

### Recent Advances in *In Silico* Tools for Expanding Reference Libraries

As can be seen from **Table 2**, measured observable data is very limited compared to the number of structures we know or suspect to exist. While many reference libraries containing experimentally determined values exist, most are currently too small or too incomplete to satisfy the needs of metabolomics studies. The most comprehensive untargeted MS-based metabolomics experiments that rely on today’s reference libraries can identify up to 10% of the observed features.^89^ However, such ratios depend on the type of data processing and assay: for GC-MS based metabolomics or in lipidomics assays, the ratio of identification is typically at 30% of features that have associated mass spectra.^90^ In fact, high quality data processing should include measures of blank sample corrections, adduct deconvolution and the use of pooled sample quality controls to reduce the number of spurious features in assessments of metabolome coverage statements.

One route for increasing the amount of observable data in reference libraries is through the synthesis or isolation of molecules of interest. However, if one assumes that the total number of all known and predicted metabolites, as well as all known anthropogenic chemicals, found in humans is ∼2 million compounds and the cost to isolate or synthesize and to comprehensively characterize these compounds is ∼$5000/chemical, such an effort would cost in excess of $10 billion USD. This initiative would easily take 20+ years and consume a significant portion of the NSF or NIH budget. In other words, the time and cost to make the comprehensive reference library required for the metabolomics community is simply not feasible. A more cost-effective approach will have to be developed. We believe that a viable option, and the future of reference library growth, is via *in silico* approaches. Simply stated, computational approaches could be used to generate *in silico* (i.e., predicted) observable data, based on validated methods. We propose this because of the foundational developments in chemistry and physics and the need to identify a vast number of unidentified features. Development of various machine learning- or quantum chemistry-based approaches (reviewed in^91^) and tools for *in silico* prediction of various types of spectra and other observables has increased the size and chemical appropriateness of existing reference libraries. Indeed, there are now several well-developed software tools for predicting electron ionization-mass spectrometry (EI-MS), electrospray ionization-tandem mass spectrometry (ESI-MS/MS) and NMR spectra, CCS, and RT values using combinatorial approaches, machine and deep learning methods, and quantum mechanical techniques. For ESI-MS/MS spectral prediction, several machine learning methods including MetFrag,^92^ CFM-ID,^93^ MS-FINDER,^94^ ChemDistiller^95^ and MAGMa^96^ have appeared. CFM-ID, MS-FINDER and MAGMa in particular have shown excellent performance in terms of spectral prediction accuracy in multiple independent tests.^97, 98^ For EI-MS spectra, two machine learning methods (CFM-ID-EI^99^ and NEIMS^100^) have been described and both perform well. Separately, a quantum mechanical method called QCEIMS^101^ has been developed to predict EI-MS spectra and more recently ESI-MS/MS spectra with QCxMS.^102^ QCEIMS and QCxMS are significantly slower than the ML methods, but they provide useful insights into the EI and ESI fragmentation processes.

Just as with EI-MS, both machine learning and quantum chemistry methods have been developed to predict NMR spectra (^1^H, ^13^C, 1D and 2D). Density Functional Theory (DFT) has been used for many years to predict NMR chemical shifts and coupling constants with errors as small as 0.2 ppm for ^1^H shifts and 2.5 ppm for ^13^C shifts.^103, 104^ This level of precision can enable distinction of closely related diastereomers.^105^ ISiCLE, which uses NWChem^106^ for calculations, is an example of a recently developed DFT method that is now being used to calculate ^1^H and ^13^C NMR spectra for thousands of natural products that do not have measured NMR spectra in NP-MRD.^82^ It is expected that ISiCLE will be able to create one of the world’s largest *in silico*-predicted NMR spectral libraries by the end of 2024. It is also possible to use machine learning and neural networks to predict the NMR spectra of small molecules.^103, 107^ These programs tend to be much faster than QM methods and may be just as accurate.^108^

Lastly, the prediction of separation properties, such as RI, RT, and CCS values, has become increasingly popular. Several machine learning-based programs for CCS prediction have recently appeared including DeepCCS,^48^ MetCCS predictor,^109^ and DarkChem.^46^ Quantum chemical methods, such as ISiCLE,^110^ have also been developed to accurately predict CCS values. Regardless of the method chosen, the typical errors between experimentally observed and predicted CCS values are as small as 2-3% with correlation coefficients greater than 0.95. Using these predictive CCS tools, several reference libraries have been generated, containing hundreds of thousands of predicted CCS values, including CCSBase,^111^ AllCCS,^47^ and MetCCS.^112^ Similar efforts are being made in RI prediction and measured data curation. The NIST 20 library contains more than 114,000 experimentally measured Kovats retention indices, used for GC-based metabolomics. Using these data, NIST scientists have recently developed a graph neural network approach that can predict retention indices with a mean absolute percentage error (MAPE) as small as 3% and a correlation coefficient of >0.98.^113^ This, by far, is the most accurate method for RI prediction ever published. A similarly accurate method for RI prediction has recently been implemented in the latest version of the HMDB which provides 6.7 million RIs for >26,000 GC-MS compatible compounds (and their derivatives).^44^ In principle, these methods could be used to generate accurate RI values for each specific GC-MS method, for hundreds of thousands of molecules which do not have experimental RI data. In terms of RT prediction for LC, several efforts aimed at relative RT prediction have been undertaken using highly specified chromatographic conditions. These include the machine learning-based tools Retip,^49^ GNN-RT,^114^ and the METLIN SMRT predictor^87^ that showed RT median prediction errors as small as 5%. However, the correlation coefficients between experimental and predicted RTs are often only ∼0.6, suggesting that RT prediction for LC has a long way to go before it matches the accuracy of RI prediction.

With increasing popularity of *in silico* approaches to generating reference observables, there have been similar efforts to predict novel metabolite structures that can be added to reference libraries. Currently, there are two approaches for doing so. One is to use enzymatic modeling to predict biotransformations or enzymatic by-products of starting molecules.^115^ The other is to use generative modeling and deep learning to create biofeasible structures.^46, 116^ Biotransformation prediction has been around for many decades and was pioneered by researchers in the drug metabolism community.^117^ As a result, a number of commercial programs have been developed, including Meteor Nexus, ADMET-Predictor, MetabolExpert and others, that predict Phase I (cytochrome P450) metabolism specifically for drug molecules and a small number of naturally occurring metabolites. These programs use expert-derived rules and large internal databases to perform look-ups and make their predictions. More recently, several open source or open access tools have appeared that perform biotransformation prediction for a larger collection of molecules. These include GLORYx,^118^ FAME 2,^119^ FAME 3,^120^ CyProduct^121^ and BioTransformer.^115^ These software packages, many of which use machine learning techniques, not only predict Phase I biotransformation, but also Phase II metabolism and microbial/gut metabolism for drugs, pesticides, herbicides and naturally occurring metabolites. Furthermore, they are also able to predict these transformations much more accurately than commercial, rules-based software.

One of these programs, BioTransformer, has recently been applied to predict the structures of 2 million biotransformed molecules (Phase I + Phase II + microbial + promiscuous enzyme transformations) using a starting set of 120,000 compounds in the HMDB. Other approaches have also made use of enzyme promiscuity to predict biofeasible metabolites. For example, the MINE (Metabolic *In silico* Network Expansions) database used an algorithm called the Biochemical Network Integrated Computational Explorer (BNICE) and expert-curated reaction rules to generate more than 570,000 biofeasible structures starting from 18,000 KEGG metabolites.^122^ At the time of writing MS-FINDER integrates structures and formulae for 224,622 known metabolites^94^ and also includes 643,307 hypothetical metabolites from MINE-DB.^122^ The advantage of these biologically based *in silico* biotransformation methods is that the enzymatic reaction steps and enzymatic mechanisms are explicitly shown or referenced. In other words, the rationale and provenance for each predicted compound is available. The disadvantage is that these biotransformation programs can occasionally produce unreasonable combinatorial explosions. Likewise, they can’t make “out-of-the-box” predictions or generate non-obvious or unexpected metabolites. An expansion of this approach was recently published to encompass likely occurring chemical damage (such as oxidations) of molecules, in an analogous database called CD-MINE to cover spontaneously occurring chemical transformations.^123^

### Guidelines for Appropriate Reference Library Size and Composition

Both the size and composition of reference libraries will impact the assessment of metabolite identification probability. A reference library that is too small can result in reduced false discovery rate and seemingly accurate, and thus overly confident, identification probabilities. One that is too large can result in increased false discovery due to the addition of compounds that are highly unlikely to be found in such a sample and reduced identification probabilities.^30^ Similarly, one should select the appropriate source of compounds to include in the reference library for a given sample type and use case. For example, if a study focuses on a specific organism in a laboratory-controlled setting, then only those molecules potentially produced or consumed by the organism, present in growth media, for example, or known as common contaminants present in the chosen analytical method should be included in the reference library. That is, to prevent misidentifications, one should use organism-specific or sample-specific reference libraries of appropriate size and composition. By comparison, the proteomics community typically uses an appropriate protein FASTA file containing the amino acid sequences of all proteins expected in the organism(s) under study and that are based on translations of the corresponding genomes when searching peptide MS/MS spectra.

### Reference libraries for studies of specific organisms in controlled laboratory settings

Different organisms can have profoundly different metabolic needs and metabolic capabilities. For instance, plants have very different metabolomes than animals.^124^ Furthermore, the regular consumption of processed foods, supplements, and drugs by humans means that people will have a very different metabolome than lab rats raised on strict chow diet. Indeed, direct comparison of plasma metabolomes showed that less than half the LC-MS signals were common to seven different mammalian species.^125^ These results argue for the need for appropriately specialized reference libraries for metabolomics studies to ensure the reliability of metabolite identification.^126^ For studies of specific organisms in controlled laboratory settings, we recommend starting with reference libraries that are populated based upon either genome-enabled metabolic reconstructions (e.g., genome-scale metabolic models)^127^ or comprehensive review and curation of the literature in respect to metabolomics and other studies of metabolisms of specific organisms. There are a number of existing genome-scale metabolic models for certain organisms, such as *E. coli*,^128^ *S. cerevisiae*,^129^ *M. musculus*, *D. rerio* and *D. melanogaster*,^130^ as well as methods and resources for the scientific community to continue to expand these models or apply them to new organisms.^131, 132^ Of note, some of these models overly depend on genomic inference and have rather sparse metabolite information, for which metabolomics can contribute significantly to their expansion.^23^ Similarly, there are a number of reference libraries for specific organisms and derived from comprehensive review and curation of the literature, such as the Yeast Metabolome Database^133^, the *E. coli* Metabolome Database^134^, and the *Pseudomonas aeruginosa* Metabolome Database.^135^ PathBank contains literature-derived metabolome reference libraries from several other model organisms.^136^ When studying specific organisms in controlled laboratory settings, our recommendation is to use a reference library derived from comprehensive review and curation of the literature. If one is not available, then use an appropriate genome-scale metabolic model. Selecting a library from another organism that is the closest taxonomical relative can also be useful in supplementing a suspect library for an organism that has not been well studied. Finally, the organism-specific metabolite reference libraries should be supplemented with additional inputs from the experiment (e.g., growth media components), including common contaminants present in the chosen analytical method and that are likely to be identified in the experimental data (e.g., plasticizers).

### Reference libraries for studies of free-living organisms or environmental systems

As mentioned above, additional considerations are necessary when developing reference libraries for free-living (e.g., humans) or environmental (e.g., soils, forests) systems. Such organisms or systems are not constrained to controlled settings and experience various and diverse inputs to their metabolomes on routine if not daily bases. Further, even within a given free-living system, a human subject for example, the metabolome of one organ or organ system can be very different than that of another. For example, the human blood metabolome is very different from the human urine metabolome.^137, 138^ Likewise, the plant leaf metabolome is very different from the associated plant rhizosphere.^139^ A number of reference libraries have been developed for free-living organisms. These include the previously mentioned HMDB,^44^ the Livestock Metabolome Database,^140^ and the Bovine Metabolome Database.^141^ Similarly, a variety of matrix-specific or biofluid-specific resources such as the Fecal Metabolome Database,^142^ the Saliva Metabolome Database,^143^ the Serum Metabolome Database,^137^ and the Urine Metabolome Database^138^ have also been publicly released. Such databases, or metabolome atlases, may also include aspects of the impact of disease or other factors. As an example, recently the Metabolome Atlas of the Aging Mouse Brain was published.^144^ As with laboratory-controlled systems, reference libraries for free-living organisms and environmental systems should be supplemented with other molecules that might be expected to be present in the organism or sample of interest, based on typical behaviors or environmental exposures. Examples of such molecules are contained within the Blood Exposome Database,^145^ which covers compounds identified in human blood; DrugBank,^146^ which covers approved drugs found in humans; the Toxic Exposome Database,^147^ which covers toxic compounds found in humans; the Norman Suspect List Exchange,^148^ which covers common environmental or water contaminants and FooDB (https://foodb.ca), which covers food compounds and food additives found in foods consumed by humans. Further, and as described above, researchers should supplement reference libraries for free-living organisms or environmental systems with information and properties for molecules that are generated via *in silico* predictions of relevant biotransformations or from *in vitro* or cellular incubations.^24^ Finally, the comprehensive reference libraries for free-living organisms and environmental systems should be supplemented with additional inputs from the experiment, including common contaminants present in the chosen analytical method and that are likely to be identified in the experimental data.

## CONCLUSIONS AND RECOMMENDATIONS FOR FUTURE IMPLEMENTATION OF METABOLITE IDENTIFICATION PROBABILITY

In this perspective, we have introduced a new concept of metabolite identification probability and have demonstrated its utility in mock identifications using reference libraries constructed from subsets of HMDB and computationally generated RT, CCS, *m/z*, and MS/MS data. The method is computationally simple, automatable, and transferable among analytical platforms. It requires only processed metabolomics data, appropriately defined tolerances allowed in the associated measurement precisions, and reference libraries that are comprehensive and appropriate for the system being queried. We recommend that the metabolomics and related communities (e.g., the non-target analysis community) join us in further exploring the metabolite identification probability approach to more fully reveal its potential and limitations, using real data from real studies and in parallel with their current preferred methods for assessing metabolite identification confidence (e.g., MSI levels), in order to accumulate data on method performance relevant to state-of-the-art. Further extension of these concepts to unidentified features will be required to fully address e.g., unknown chemical hazards of the exposome.^64, 149^

Metabolite identification probability is heavily dependent on the richness of the experimental data being matched to the reference library, the dimensionality and therefore overall resolution of the analytical measurement, the overall measurement precision(s), and the composition and size of the reference library itself. A key requirement for successful implementation of the metabolite identification probability concept is thus the availability of comprehensive and system-appropriate reference libraries. Further research and discussion within the community are needed to determine the repertoire of metabolites and related molecules that should comprise a reference library for a given system, such that metabolite identification probabilities are neither over- nor underestimated. Related, because of the limitation of commercial availability of reference compounds for all system-relevant small molecules, we recommend that the community begin adopting computational approaches for calculating or predicting the associated observable properties, such as spectra, such that reference libraries can be made complete. The accuracy of computationally predicted data should improve with time as methods and technology improve.

Finally, in order that reported metabolite identification probabilities can be transparent, we recommend that individual laboratories version their in-house reference libraries and make them available to the rest of the community as e.g., open mass spectral libraries (OMSL). Besides increasing transparency in calculations of identification probabilities, versioned OMSL and other libraries will be a tremendous resource to the metabolomics research community, as has already been demonstrated by resources such as GNPS^79^ and enabled through workflows such as FragHub.^150^ As inspiration for how such sharing might be implemented, the metabolomics community can look to the Universal Protein Knowledgebase (UniProtKB)^151^ as an example. UniProtKB is a freely accessible database of curated protein sequences that are used, among other purposes, as “reference libraries” for proteomics data searches.

## ACKNOWLEDGMENTS

The concept of metabolite identification probability emanated from discussions held among the Compound Identification Development Cores Sub-Committee of the NIH Common Fund Metabolomics Program Phase II. The authors thank Mr. Nathan Johnson from Pacific Northwest National Laboratory (PNNL) for help in designing Figure 1. T.O.M., C.H.C, V.G., A.A., S.T., F.W. S.M.C., J.R.N., M.R.B., M.R.S., and D.S.W. were supported by the National Institutes of Health, National Institute of Environmental Health Sciences (NIEHS) grant U2CES030170 via the Pacific Northwest Advanced Compound Identification Core. A.S.E. was supported by NIEHS grant U2CES030167. D.P.J. and E.T.M. acknowledge support from NIEHS grant U2CES030163. D.S.W. acknowledges additional support from the Natural Sciences and Engineering Research Council of Canada (NSERC) and from the Canada Foundation for Innovation Major Science Initiative (CFI-MSI). G.J.P. acknowledges support from National Cancer Institute grant U01CA235482 and NIEHS grant R35ES028365. O.F. was supported by NIEHS grant U2CES030158 and National Institute of General Medical Sciences grant R01GM0155383. T.O.M. and D.H.R. acknowledge additional support from the PNNL Laboratory Directed Research and Development Program via the *m/q* Initiative.

PNNL is a multi-program national laboratory operated by Battelle for the U.S. Department of Energy under Contract DE-AC05-76RLO 1830.

This manuscript was subjected to the U.S. Environmental Protection Agency internal review process. The research results presented do not necessarily reflect the views of the Agency or its policy. Mention of trade names or products does not constitute endorsement or recommendation for use.

