## Supplemental Figures for "Introducing ‘identification probability’ for automated and transferable assessment of metabolite identification confidence in metabolomics and related studies"

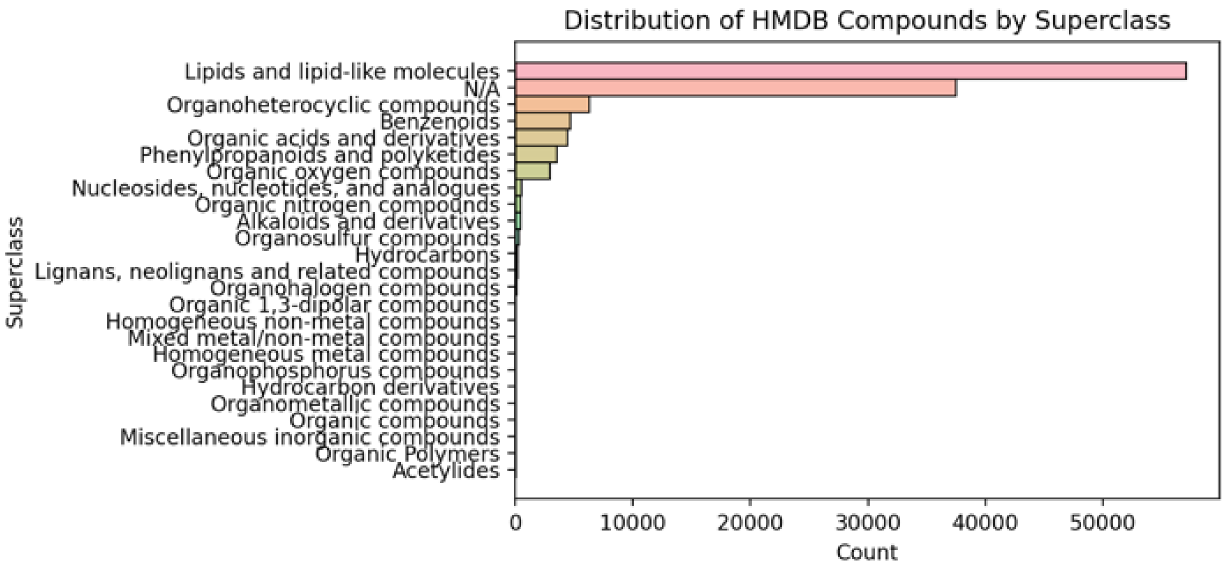


**Supplemental Figure S1. Distribution of ClassyFire superclass annotations for all compounds contained in HMDB, accessed July 14, 2022.** Note that he “nonlipids” designation excludes compounds categorized into the “Lipids and lipid-like molecules” and “N/A” superclasses.
